# Data-Driven Phenotypic Categorization for Neurobiological Analyses: Beyond DSM-5 Labels

**DOI:** 10.1101/051789

**Authors:** Nicholas T. Van Dam, David O’Connor, Enitan T. Marcelle, Erica J. Ho, R. Cameron Craddock, Russell H. Tobe, Vilma Gabbay, James J. Hudziak, F. Xavier Castellanos, Bennett L. Leventhal, Michael P. Milham

## Abstract

**Background:** Data-driven approaches can capture behavioral and biological variation currently unaccounted for by contemporary diagnostic categories, thereby enhancing the ability of neurobiological studies to characterize brain-behavior relationships.

**Methods:** A community-ascertained sample of individuals (N=347, ages 18–59) completed a battery of behavioral measures, psychiatric assessment, and resting state functional magnetic resonance imaging (R-fMRI) in a cross-sectional design. Bootstrap-based exploratory factor analysis was applied to 49 phenotypic subscales from 10 measures. Hybrid Hierarchical Clustering was applied to resultant factor scores to identify nested groups. Adjacent groups were compared via independent samples t-tests and chi-square tests of factor scores, syndrome scores, and psychiatric prevalence. Multivariate Distance Matrix Regression examined functional connectome differences between adjacent groups.

**Results:** Reduction yielded six factors, which explained 77.8% and 65.4% of the variance in exploratory and constrained exploratory models, respectively. Hybrid Hierarchical Clustering of these 6 factors identified 2, 4, and 8 nested groups (i.e., phenotypic communities). At the highest clustering level, the algorithm differentiated functionally adaptive and maladaptive groups. At the middle clustering level, groups were separated by problem type (maladaptive groups; internalizing vs. externalizing problems) and behavioral type (adaptive groups; sensation-seeking vs. extraverted/emotionally stable). Unique phenotypic profiles were also evident at the lowest clustering level. Group comparisons exhibited significant differences in intrinsic functional connectivity at the highest clustering level in somatomotor, thalamic, basal ganglia, and limbic networks.

**Conclusions:** Data-driven approaches for identifying homogenous subgroups, spanning typical function to dysfunction not only yielded clinically meaningful groups, but captured behavioral and neurobiological variation among healthy individuals as well.

## Introduction

The limitations of categorical definitions of psychiatric illness for clinical practice (1) and psychiatric research (2) are increasingly apparent. While diagnostic labels defined in nosological systems such as the Diagnostic and Statistical Manual of Mental Disorders (DSM (3)) are needed for clinical practice, these systems impede the search for pathophysiological markers using epidemiologic, genetic, and neuroimaging approaches (4). Given growing recognition of these limitations, the Research Domain Criteria (RDoC) Project has called for the development of a new nosology (5). In response, empirical data are being used to identify target phenotypic domains and constructs to characterize psychopathology and guide psychological and neurobiological investigations.

Not surprisingly, how to best delineate phenotypic domains/constructs to guide a non-syndromal research framework is uncertain (6). Inherent to this pursuit is the varying utility of categorical and dimensional frameworks. While dimensional models of psychopathology are widely supported (7–9), fully dimensional perspectives have limitations with respect to clinical decision-making (10, 11). Further, it is unclear whether it is more expedient to derive phenotypic targets from existing models (based on existing data and theory), data-driven analytic approaches (12, 13), or some combination. Psychiatric classification systems have variable derivations spanning clinical/research observations (e.g., DSM) to empirical assessment (e.g., Achenbach System), with other entities exhibiting a combination (e.g., RDoC). Consensus-driven methods (e.g., DSM; also arguably RDoC – see (14)) can certainly provide valuable insights; however, data-driven approaches may be crucial for identifying more behaviorally refined biological phenotypes (15) to address the profound heterogeneity evident in health and illness (16, 17). Fair, Nigg, and colleagues recently demonstrated the potential value of delineating groups by similarity/dissimilarity of individual phenotypic profiles (e.g., neuropsychological profiles, temperament profiles). Adopting community detection methodologies from graph theory, they successfully identified: i) six distinct neuropsychological profiles that capture normal variation and are modified by Attention-Deficit/Hyperactivity Disorder (18) and ii) three temperamental phenotypes that showed intriguing biological differences as well as differential clinical outcomes (19).

Here, we used the Nathan Kline Institute Rockland Sample (NKI-RS)(20), a deeply phenotyped, community-ascertained multimodal imaging sample. Using data from adult participants (ages 18–59) in the NKI-RS, we aimed to identify data-driven phenotypes, based on core behavioral features representing several domains of function (including personality/temperament, symptom features, interpersonal functioning, and behavioral tendencies). Our first aim is to identify phenotypic dimensions that accurately represent meaningful variation across multiple domains of behavior. Accordingly, we conducted a bootstrap-based exploratory factor analysis on 49 subscales derived from 10 measures obtained for 347 participants. The second aim is to identify a nested hierarchy of homogenous participant groups via hybrid hierarchical clustering of participants, based on the factor profiles that we previously identified. To provide a phenotypic characterization of the participant groupings identified, we used DSM-IV labels and Achenbach’s Adult Self-Report (ASR (21)) – neither of which were included in the factor analysis. The third aim is to examine multivariate intrinsic brain functional connectivity differences among adjacent clusters/groups (derived from the first two aims).

## Materials and Methods

### Participants

Participants were recruited as part of the NKI-RS (20), a community-based sample of approximately 1,000 participants, ages 6-85 years. To ascertain a cohort approximating a representative sample, exclusion criteria were minimal. Notably, comorbid medical conditions and medications (including psychotropics) were permitted. Written informed consent was obtained from all participants in accordance with local Institutional Review Board oversight. The following inclusion criteria were applied: (i) ages 18-59; (ii) absence of serious head injury and/or major neurological disorder; (iii) negative history of bipolar disorder or psychosis; (iv) negative drug test for commonly used illicit drugs with no therapeutic analogs/applications; (v) at least 95% completion of each self-report measure examined.

### Subject Phenotyping

All participants completed the Structured Clinical Interview for the DSM (SCID (22)), and the Edinburgh Handedness Inventory (EHI (23)), in addition to measures reflecting clinical symptom domains, personality/temperament, and broad behavioral characteristics (see **Table S1**; also (20)). We selected subscales rather than individual items or full-scale scores to balance depth of assessment and amount of data per subject.

### MRI Acquisition

Imaging data were collected on a 3T Siemens TIM Trio system equipped with 32-channel head coil. Both structural and resting state functional magnetic imaging (R-fMRI) data were acquired. The structural image was a T1-weighted magnetization prepared gradient echo sequence (MPRAGE): TR=1900 ms, TE=2.52 ms, flip angle=9°, 176 slices, 1 mm^3^ isotropic voxels. The R-fMRI data were acquired via multiband echo-planar imaging (MB-EPI(24)) with the following parameters: volumes=900, TR=645 ms, TE=30 ms, flip angle=60°, 3mm^3^ isotropic voxels.

### Phenotypic Analysis

#### Data Screening

All self-report data were checked for univariate and multivariate outliers. We also tested the assumption of missingness at random (MAR(25)). Missing data were imputed using an expectation-maximization algorithm (26).

#### Dimension Reduction

For each participant, we first created a multidimensional phenotypic profile using 49 subscale scores obtained from 10 different questionnaires. Given modest intercorrelations among the differing questionnaires, we next performed an exploratory factor analysis to obtain a reduced set of dimensions. Analyses were done on age-and gender-regressed residuals of the 49 subscale scores to minimize the impact of these demographic variables on clustering (27). Parallel analysis (28) of 10,000 permutations of the raw data (29), was used to determine number of factors, comparing raw data eigenvalues to those obtained in the 95^th^ percentile of the permutations. Maximum likelihood factor estimation with varimax rotation was used to estimate six factor loadings for each subscale score. Confidence intervals for factor loadings were estimated from 10,000 bootstrapped re-samplings (30). To minimize factor intercorrelations, factor loadings overlapping zero (95% CI) and/or exhibiting values < 0.25 were set to 0 in a restricted model. Factor scores (i.e., equivalent of latent value per factor) were computed using regression estimation (31).

#### Clustering Analysis

Categorical approaches (e.g., DSM) confer the ability to delineate unique combinations of above-threshold impairments in subsets of individuals. Building on this strength, a central goal of the present work is to identify phenotypically distinct groupings of participants among the larger sample (akin to categories), based on their dimensional 6-factor phenotypic profiles. Specifically, we implemented hybrid hierarchical clustering (HHC)(32) using tree-structured vector quantization (33) to identify nested participant groups based on Euclidean distances between participant factor score profiles. HHC combines agglomerative and divisive clustering by (i) identifying mutual clusters (i.e., groups of data that are exceptionally close to one another and as a group, distant from all others) via agglomerative clustering, (ii) implementing constrained divisive clustering (retaining mutual clusters), and (iii) applying additional divisive clustering, which explores the division of mutual clusters. In combination with visual examination of the dendrogram, we used the Calinski-Harabasz criterion (CHC)(34) at each cluster number to inform cut decisions (i.e., where to divide into subgroups).

#### Cluster comparisons

Pair-wise comparisons were made among adjacent clusters at each level using EFA, ASR (21), and SCID profiles. EFA scores were fully dimensional, while the ASR and SCID diagnoses were categorical. A categorical version of ASR scores was achieved by mimicking common clinical strategies for its use, which apply cut-scores to identify meaningful psychopathology. For each grouping of participants identified via cluster analysis, we calculated the percentage of individuals exhibiting standardized scores (*T*-scores) ≥ 60 within each ASR domain. The cutoff of 60 was chosen to increase sensitivity to subthreshold symptoms of potential relevance. EFA scores were compared using independent sample t-tests, while ASR proportions and Psychiatric Diagnoses were compared using chi-square tests. We also conducted t-tests on continuous ASR scores, reported in **Supplemental Information**.

### MRI Data Processing

#### R-ƒMRI Preprocessing

Data were preprocessed using the Configurable Pipeline for Analysis of Connectomes (C-PAC; http://fcp-indi.github.io), which combines tools from AFNI (http://afni.nimh.nih.gov/afni), FSL (http://www.fmrib.ox.ac.uk), and Advanced Normalization Tools (ANTs; http://stnava.github.io/ANTs), using Nipype (35). Preprocessing included: (i) motion correction, (ii) mean-based intensity normalization, (iii) nuisance signal regression, (iv) temporal band-pass filtering (0.01-0.1 Hz), (v) co-registration of functional to structural images using boundary-based registration (36) using FSL’s FLIRT (37), (vi) normalization of functional to MNI152 template (38) by applying a nonlinear transform from ANTs, and (vii) smoothing with a full-width-at-half-maximum 6mm Gaussian kernel. Nuisance regression removed linear and quadratic trends to account for scanner drift, 24 motion parameters, and 5 “nuisance” signals, identified via the component correction approach (CompCor (39)).

#### Multivariate Distance Matrix Regression (MDMR)

At each level of the hierarchy identified via HHC, we used MDMR to compare voxel-wise functional connectivity profiles between adjacent phenotypic groups (e.g., C1 vs. C2, C1a vs. C1b, C2a vs. C2b).

MDMR was performed on a voxel-by-voxel basis. At each voxel, the following three steps were carried out: (i) for each participant, Pearson’s correlations were computed between the target voxel and all other voxels within a specified brain mask; this step generated, for each participant, a whole-brain functional connectivity for the target voxel; (ii) a between-participant distance matrix was computed, in which each entry is the distance between the connectivity maps obtained for the target voxel in two different participants. Distance is defined as √(2 * (1 – *r*)), where *r* is the spatial correlation of the connectivity maps obtained at the target voxel in two different individuals. Importantly, these distances are calculated independent of any phenotypic relationships; (iii) a pseudo-*F* statistic was computed to provide mathematical evaluation of the relationship between the variability in the distance matrix (40) computed in step ii and the variable of interest (i.e., group membership). Thus the pseudo-*F* value at each voxel tells us whether the functional connectivity profiles for that voxel varied among individuals as a function of the phenotypic group membership (e.g., C1 vs. C2, C1a vs. C1b, C2a vs. C2b).

MDMR was applied using the ‘Connectir’ package in R (http://czarrar.github.io/connectir) on resampled, 4mm^3^ isotropic voxels. Computations were constrained to a study-specific group mask, including only voxels present across all participants and contained in a 25% probability gray-matter MNI mask. The MDMR model (at each level of the dendrogram) specified cluster membership (categorical) and age, sex, hand laterality, and mean framewise displacement (FD) as covariates.

Consistent with prior work, voxel-wise significance of the pseudo-*F* statistic was determined via estimation of the null distribution with random permutation (n=10,000) for each cluster/community comparison (40). Recent work has raised concerns about potentially inflated type I error rates with Random Field Theory (RFT) cluster thresholding approaches (cf. (41, 42)). These concerns are primarily applicable to parametric tests; non-parametric approaches are likely less prone to such inflations.

However, we have chosen a more conservative thresholding approach than those used in prior studies implementing MDMR. Specifically, we corrected for multiple comparisons using cluster-based permutation (n=5000) with a height threshold of *Z* ≥ 2.33 (*p* < 0.01) and cluster extent probability of *p* < 0.05.

Given that lower levels of the hierarchy had few participants per group, statistical power in these comparisons would be expected to be notably lower. For illustrative purposes, we repeated analyses using less stringent criteria (voxel-wise p < 0.05; RFT-corrected p < 0.05) and included these results in the **Supplementary Information**.

## Results

Demographic and diagnostic information is provided in **Table 1**. Details about data screening are provided in the **Supplementary Information**.

**Table 1.**
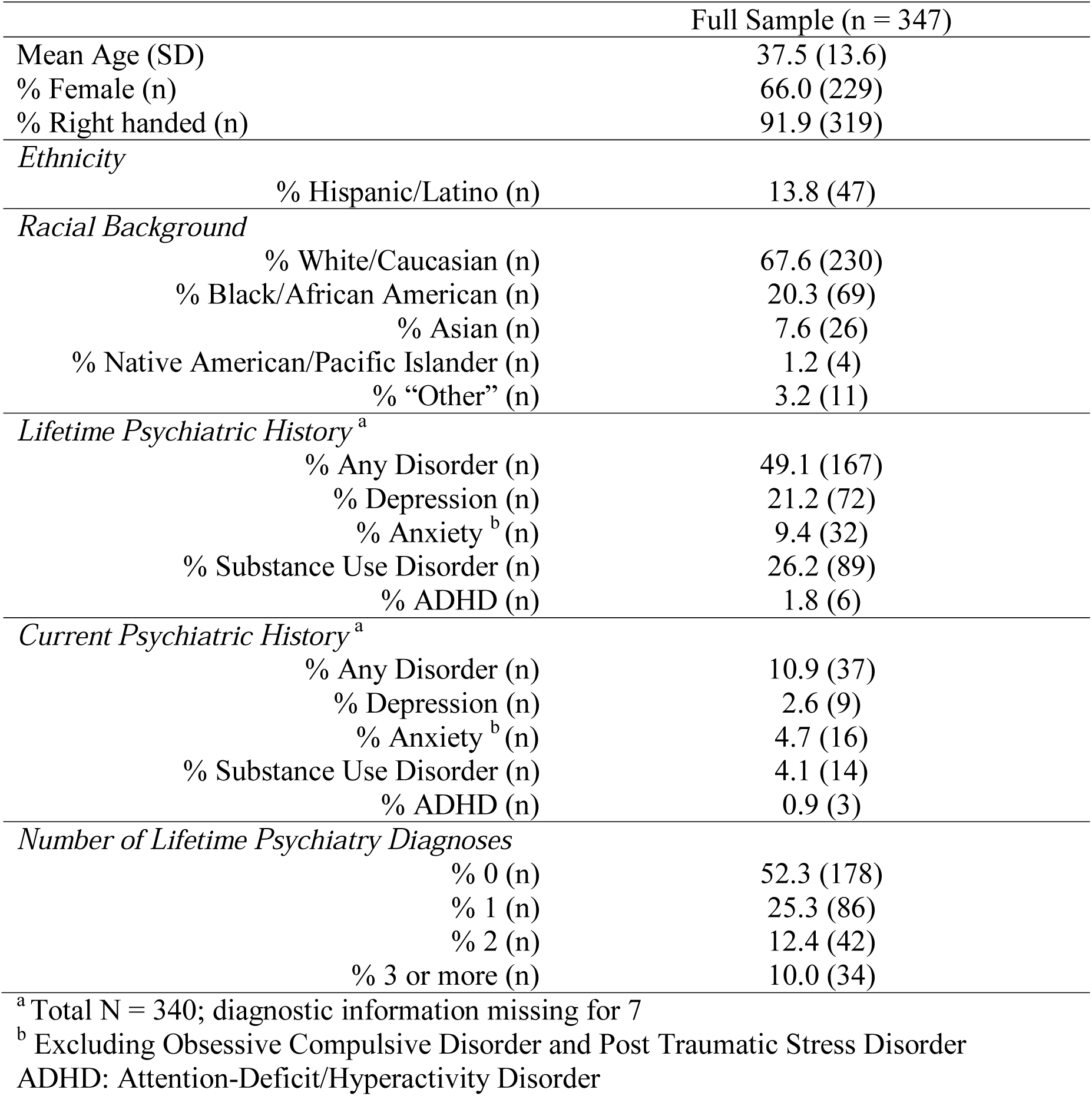
Demographic and Diagnostic Characteristics

### Dimension Reduction

A 6-factor solution was estimated with maximum likelihood-based exploratory factor analysis, accounting for 77.8% of the variance. The constrained model (eliminating low-loading items) accounted for 65.4% of the total variance. Standardized factor loadings are presented in **Table S3**. Correlations between latent factor scores and regression estimates are displayed in **Table S4**. Primary factor loadings and example items for each subscale are provided in **Figure 1**. The 6 retained factors were interpreted as follows (see **Figure 1**): (i) General Distress & Impairment, (ii) Conscientiousness, (iii) Sensation & Risk Seeking, (iv) Frustration Intolerance, (v) Contextual Sensitivity, and (vi) Neuroticism & Negative Affect. Details about each of the factors are provided in **Supplementary Information**.

**Figure 1.**
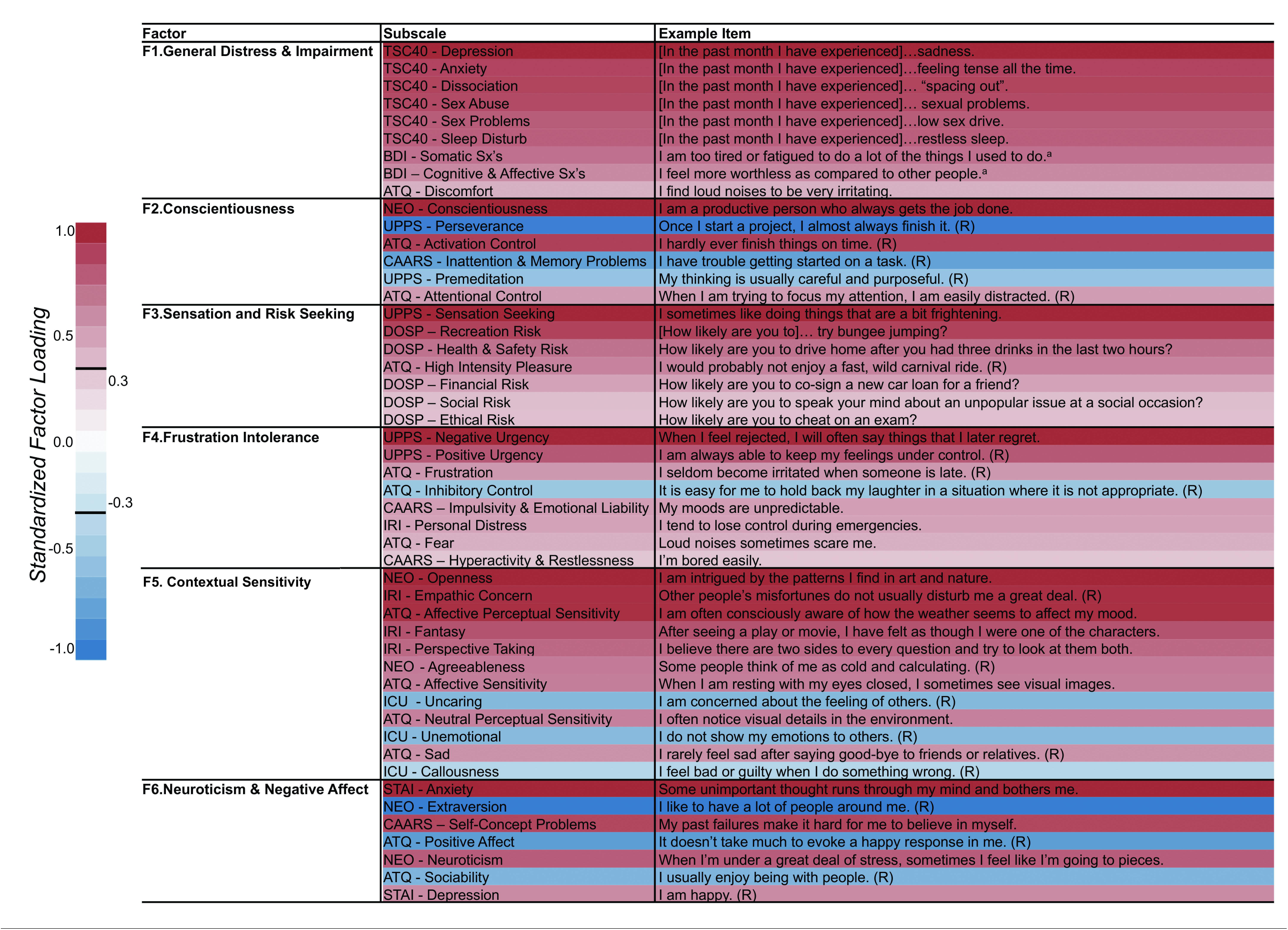
Factors identified by bootstrap-based maximum likelihood exploratory factor analysis. Factors and their corresponding subscales (along with example items) identified by exploratory analysis with 10,000 bootstrap resamplings. Subscale names are provided in the middle column. Sample items for each subscale are provided in the far right column. Color bar on the left provides an index to the shading of each subscale relative to its standardized loading on the factor. ATQ = Adult Temperament Questionnaire; BDI = Beck Depression Inventory, 2^nd^ Edition; CAARS = Conners’ Adult ADHD Rating Scale; DOSP = Domain-Specific Risk-Taking Scale; NEO = NEO - Five Factor Inventory; ICU = Inventory of Callous and Unemotional Traits; IRI = Interpersonal Reactivity Index; STAI = Spielberger State Trait Anxiety Inventory; TSC40 = Trauma Symptom Checklist; UPPS = Impulsive Behavior Scale. Notes: (R) = reverse-scored item in scale. ^a^ These sample items were selected from the response option corresponding to 2 on a 0-3 scale.

### Cluster Analysis

Visual inspection of the dendrogram suggested three clear cut-points, yielding 2, 4, and 8 groups, respectively. The largest CHC value was observed at *k*=2 and stable subgroups (operationalized as values of *k* wherein CHC did not change appreciably from the prior solution (i.e., local minima)) at *k*=4 and *k*=8. Visual examination of participant-by-participant correlation and squared Euclidean distance matrices supported the face validity of these cut-points (see **Figure 2**).

**Figure 2.**
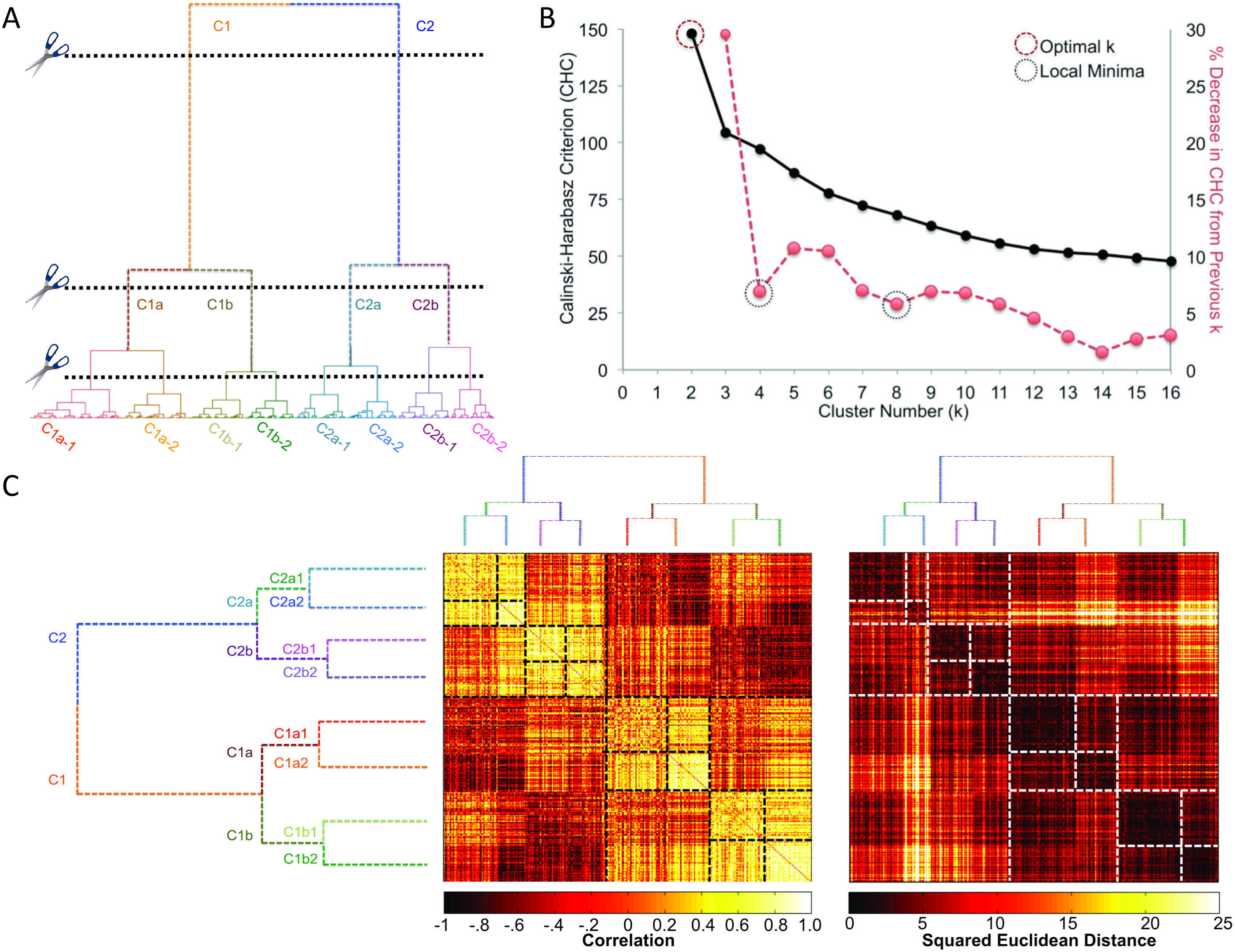
Results of hierarchical clustering. Panels in this figure depict the decision-criteria used to ascertain clustering levels and resultant groups, as well as the similarity and dissimilarity of the groups as a function of correlation and squared Euclidean distance between factor scores (subject-by-subject). **Panel A** shows the dendrogram resulting from Hybrid Hierarchical Clustering. It also shows the various levels at which the dendrogram was cut and the resultant groups. **Panel B** shows the Calinski-Harabasz Criterion (CHC; black line), used as a decision aid for dendrogram cutting, as a function of cluster number. The red line depicts percent change in the CHC value from one cluster to the next. Since CHC did not exhibit a typical pattern (i.e., elevation at some cluster level), we defined stability (i.e., minimal change from one cluster number to the next) as our goal in deciding where to cut the dendrogram. **Panel C** again depicts the dendrogram, but relative to the correlation matrix and squared Euclidean distance matrix. Groups and subgroups are outlined with dashed lines to help visualize group membership and increased similarity/decreased dissimilarity.

### Phenotypic Cluster Differences

At all 3 levels of the dendrogram (i.e., 2-, 4- and 8- cluster solutions), the participant groups differed from one another with respect to their phenotypic profiles. Due to space limitations, we limit reported findings beyond the first level (C1 vs. C2) to those along the C2 arm (which exhibited more psychopathology-like patterns). Phenotypic results along the C1 arm, as well as for both groups together are provided in **Supplementary Information**.

#### Level 1

Results at level 1 were robust, reflecting broad-reaching group differences that spanned nearly all domains. More specifically, cluster 1 (C1) participants exhibited higher levels of adaptive functionality and cluster 2 (C2) higher levels of maladaptive functionality; significant differences were noted in nearly all measures included in the three phenotypic profiles examined (Exploratory Factor Analysis [EFA], Achenbach Adult Self-Report [ASR], Psychiatric Diagnosis [SCID]) (see **Figures 3**, **S1**). To facilitate comparison, we also depicted phenotypic findings as heatmaps (and using a continuous score on the ASR) in **Figure S2**.

**Figure 3.**
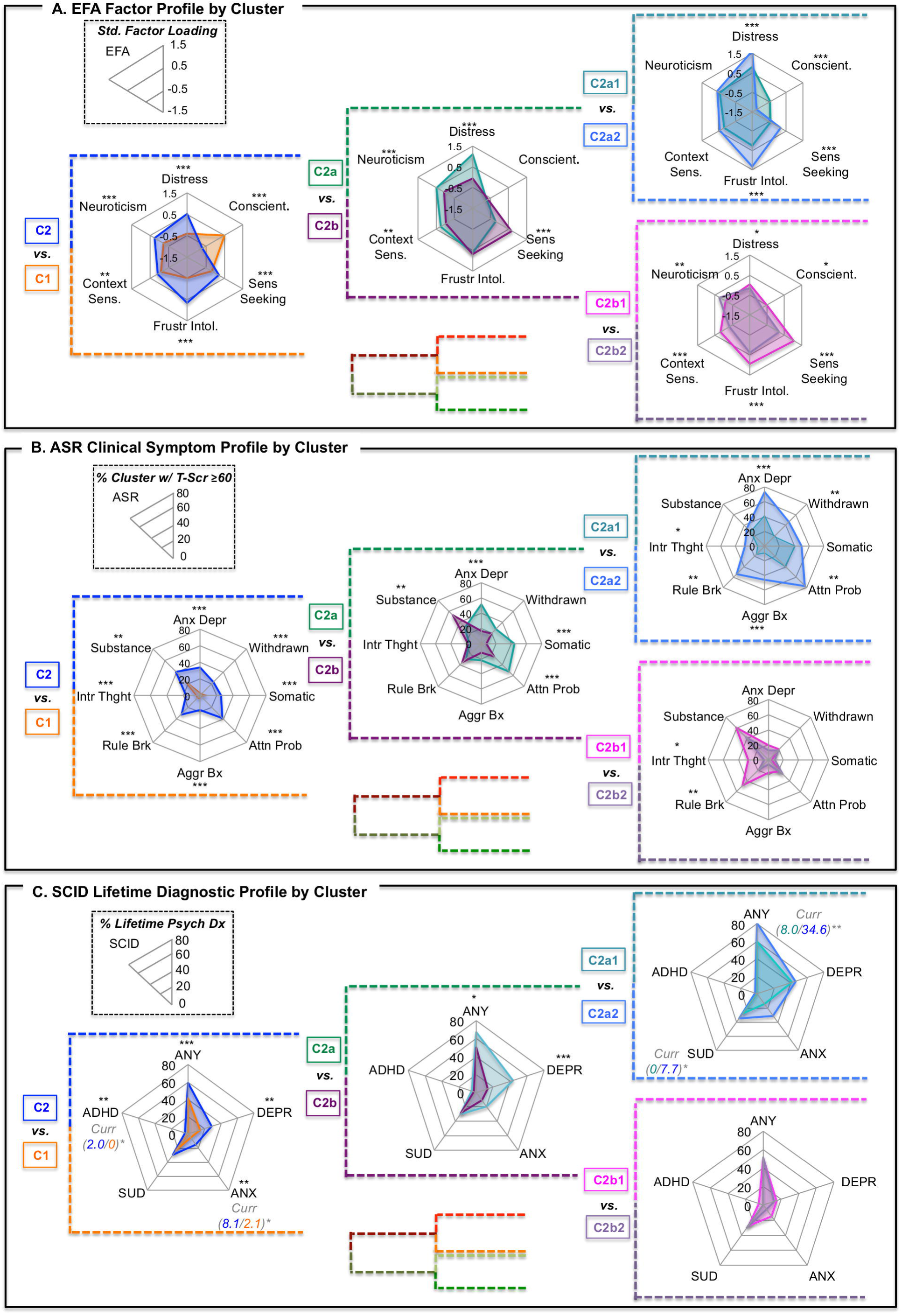
Factor, clinical symptom, and lifetime psychiatric profiles visualized as radar plots by cluster/group at three levels of hierarchical clustering, showing expansion within C2. All panels in this figure represent different measures pertaining to the same clusters (i.e., C2 is the same group of individuals, showing variation in factor profiles – Panel A; ASR clinical symptom profiles – Panel B; and lifetime psychiatric diagnosis – Panel C). **Panel A** represents mean values by cluster for each of the 6 factors from the exploratory factor analysis. Plots represent a standard loading of −1.5 at the origin and 1.5 at the maximum for each of the 6 factors. **Panel B** represents percent of individuals within a cluster exhibiting *T-*scores ≥ 60 (1 standard deviation above the mean; approaching clinical importance) for 8 domains from the Achenbach Adult Self-Report. Plots represent 0 at the center and 25% at the periphery (unless otherwise denoted) for each of the 8 domains. **Panel C** represents percent of individuals within a cluster exhibiting a lifetime (i.e., past or current) psychiatric diagnosis (ANY = any diagnosis; DEPR = depressive disorder; ANX = anxiety disorder, excluding OCD and PTSD; SUD = substance use disorder; ADHD = attention-deficit/hyperactivity disorder). Where significant differences in current psychiatric diagnosis were observed, group percentages and significance is represented next to the diagnosis in italics. Plots represent 0 at the center and 60% at the periphery (unless otherwise denoted) for each of the 5 diagnostic categories. Note that diagnoses are not mutually exclusive. Significant group differences are represented by asterisks; * *p* < .05, ** *p* < .01, *** *p* < .001.

#### Level 2

The second level (k=4) subdivided C1 (C1a and C1b) and C2 (C2a and C2b). C2 (functionally maladaptive group) was further divided into internalizing (C2a) and externalizing (C2b) problem characteristics. C2a had significantly higher ASR scores across all internalizing domains (see **Fig S2**). C2a also exhibited significantly higher rates of any lifetime psychiatric diagnosis and lifetime depression. C2b exhibited significantly higher levels of Sensation and Risk Seeking on the EFA Factor Profile and significantly higher levels of ASR externalizing problems (see **Figure 3**).

#### Level 3

The third level (k=8) divided the 4 clusters from *level 2* into 8 total sub-clusters (two clusters at level 3 for each cluster at level 2). Significant pairwise differences in ASR domains were largely a difference of magnitude (see **Figures 3**, **S1, S2**). Overall there were few significant pairwise differences between sub-clusters in DSM diagnoses, though notably C2a2 exhibited more current psychopathology than C2a1.

### Multivariate Intrinsic Connectivity Differences Among Clusters

With permutation-based cluster correction, only MDMR findings from the first level (C1 vs. C2) survived multiple comparison correction (see **Figure 4;** see **Table S8** for functional peaks). This is not surprising given the larger number of participants in each group at the first level (C1-functionally adaptive: n = 165; C2-functionally maladaptive: n =115), compared to the lower levels, which subdivided the sample into 4 and 8 groups, respectively.

**Figure 4.**
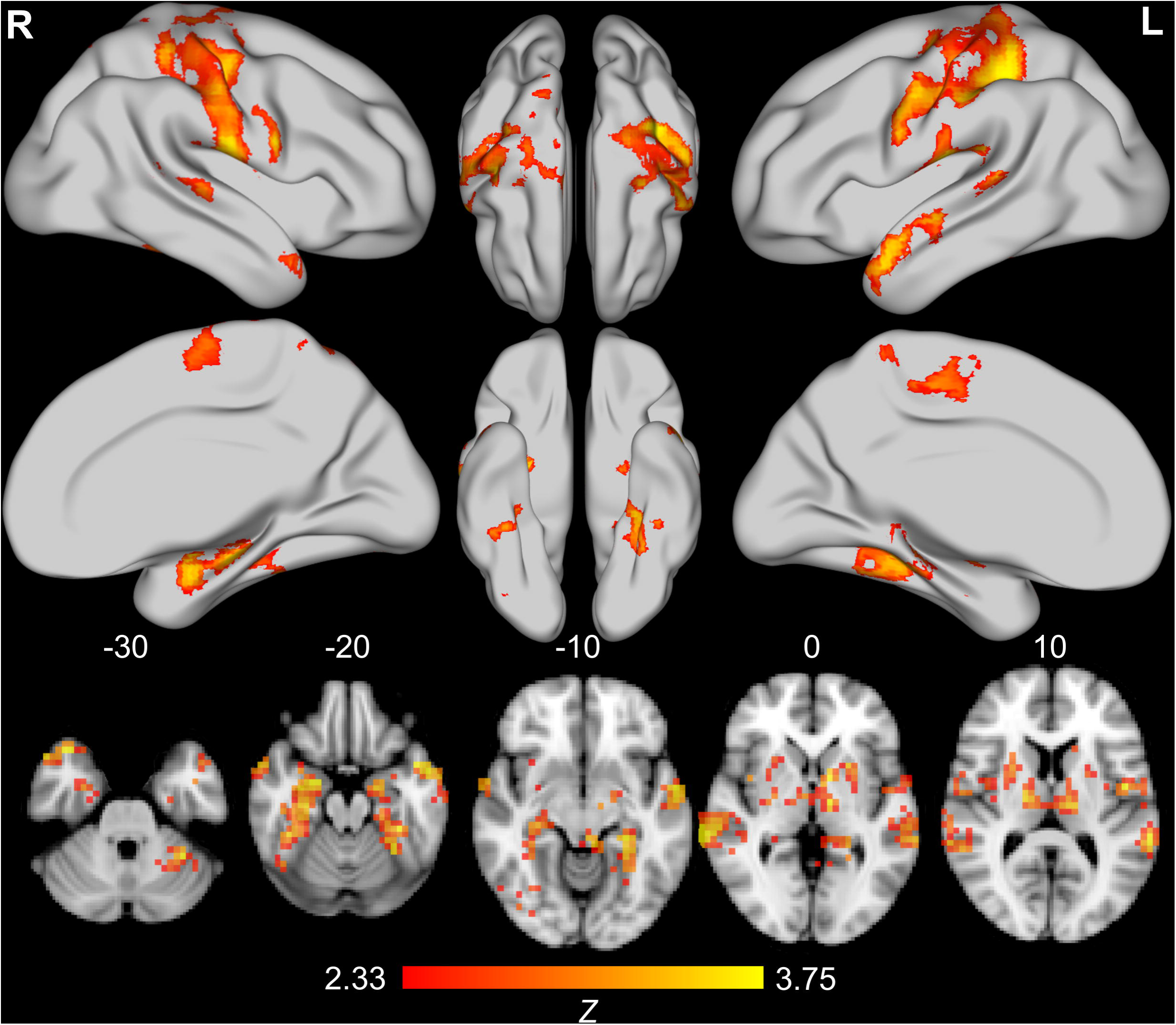
Results from multivariate distance matrix analysis of the functional connectome between C1 and C2. Adjacent groups at the highest level of hierarchical clustering (level 1: n_C1_=165, n_C2_=115) are displayed. Rendered brains and axial slices reflect multivariate distance matrix regression comparing intrinsic connectivity between groups; findings represent conversion of pseudo-*F* test results to Z values via permutation testing (10,000 resamplings of data) and permutation-based cluster correction (5,000 resamplings of data) with cluster formation set at *p* < .01 and extent threshold set at *p* < .05. Note that images are presented in neurological convention (L=R, R=L). The bottom row of axial slices represents significant findings at MNI axis Z values of −30, −20, −10, 0, and 10. From left to right, top to bottom, the top 2/3 of the figure depicts the lateral and medial surface of the right hemisphere, the dorsal and ventral surface of the right, then left, hemisphere, and finally, the dorsal and medial surface of the left hemisphere. Only Level 1 results survived cluster permutation testing correction for multiple comparisons.

Three clusters were identified for the first level comparison of C1 vs. C2. The largest cluster (*k*=133,248mm^3^, *p*=.0005) included the bilateral primary and secondary somatosensory cortices, as well as premotor, motor, and supplementary motor regions and was approximately centered on the midline near the supplementary motor area (*X*=0, *Y*=-20, *Z*=54); it also extended bilaterally to the lateral temporal lobes. These regions have been implicated in bodily self-awareness (43) and interoception (44), and identified as comprising a major hub (45) and critical network (46) of the functional brain. Additionally, emerging evidence suggests an important role for the somatosensory/somatomotor hub in high prevalence psychopathology (e.g., (47, 48)).

The second largest cluster (*k*=27,323mm^3^, *p=*.0061) was approximately centered on the left thalamus (*X*=-16, *Y*=-20, *Z*=14). In addition to a large thalamic contribution, the cluster included limbic regions (e.g., hippocampus, amygdala), decision-making regions (e.g., caudate, putamen), and various language (e.g., lingual gyrus) and vision regions (e.g., fusiform gyrus). It also extended from the left to right thalamus into the right caudate and putamen. The cluster comprised thalamic and basal ganglia regions commonly implicated in models of mental illness that emphasize thalamocortical and frontostriatal contributions (e.g., (48, 49))

Finally, the third cluster (*k=*14,528mm^3^, *p=*.0167) was approximately centered on the right hippocampus (*X*=32, *Y*=-24, *Z*=-18) and was largely comprised of right limbic regions (e.g., hippocampus and amygdala), as well as the caudate/putamen, fusiform gyrus, and middle and posterior insula. The regions implicated in this cluster, especially the right hippocampus and amygdala, are commonly associated with automated emotional processing (e.g., (50)), particularly of the kind related to high prevalence psychopathological alterations (e.g., (51)). Regions herein have also been identified as part of the medial temporal lobe subsystem of the default mode network (DMN)(52).

See supplementary materials for MDMR findings obtained at the 2^nd^ and 3^rd^ level of the dendrogram using a more liberal thresholding strategy.

## Discussion

Traditional psychiatric nosology comprises heterogeneous categories that exhibit few meaningful neurobiological correlates. The present findings illustrate the utility of data-driven approaches to: (i) derive relevant phenotypic dimensions from diverse measures, (ii) identify interpretable groups from hierarchical clustering of dimensional variables, and (iii) use nested groups as a basis of comparison for neurobiological measures.

Our findings suggest that the inclusion of instruments with normal distributions (i.e., not truncated due to an assessment floor, such as an absence of symptoms; e.g., the NEO-FFI, ATQ) can be critical to defining relevant groups in population-based classification and among high-prevalence conditions (e.g., anxiety, depression, substance use). While most individuals in the functionally adaptive group had truncated scores on syndrome-focused or problem-focused measures (somewhat remedied by using raw symptom scores), our inclusion of bipolar scales allowed us to delineate further subgroups. The present work also highlights the impact of including assessment tools that capture positive or protective factors rather than merely focusing on syndrome characteristics or problems. For example, conscientiousness, a personality characteristic associated with health and well-being (53), was a large contributor to differentiating groups at the highest and lowest levels of the nested hierarchical classification. Protective factors are rarely assessed in syndrome-or problem-focused assessments, though they can have important implications for presentation and prognosis among neuropsychiatric conditions (54).

Within the limitations of the current sample size, the present work demonstrated the value of pursuing nested subgroups within both the functionally maladaptive (C2) and functionally adaptive (C1) participant groupings. Of concern, it is possible that the groups merely recapitulated the theories on which some of the measures were predicated. It is important to note, however, that no single scale had its sub-scores distributed in a manner that could explain all components derived from the factor analysis (arguably, the NEO-FFI came the closest). We observed some truly novel combinations of subscales in the factors derived, reflecting a range of psychiatric symptoms, personality features, and protective/risk factors.

The present data also show the value of using broad behavioral characteristics to identify groups for the purposes of exploring potential neurobiological differences. Multivariate comparisons of intrinsic connectivity differences were implemented at three different clustering levels, though the findings only passed stringent multiple comparisons correction at the first level, where power was the largest. The connectome differences observed between the functionally maladaptive and adaptive groups (C1 vs. C2) at the first level were evident within (i) the somatomotor network, (ii) thalamic and basal ganglia regions, and (iii) the amygdala and extended hippocampal complex.

The somatomotor network is a somewhat novel functional target in the context of identifying potential imaging biomarkers of the tendency towards psychiatric illness. While known as a key hub (45, 46) of the functional brain, the potential role in psychopathology of the somatomotor network is only beginning to be recognized (47, 48). The somato-sensory/motor network is intimately involved in bodily self-consciousness and interoception (43, 44), processes that are increasingly implicated in predictive outcome models (55), especially for neuropsychiatric illness (56). At a coarse level of group differentiation (e.g., functionally maladaptive vs. functionally adaptive), these results underscore the potential importance of subjective valuation and bodily states in making interactional predictions that may be fundamentally altered as part of the pathophysiology of psychiatric illness.

Somewhat less novel, connectomic alterations in the thalamus and basal ganglia, as well as the amygdala and extended hippocampal complex, underscore the importance of basic cognitive functions (e.g., attention, working memory), as well as reward and emotion/saliency in delineating adaptive from maladaptive function. This latter finding lends support to the RDoC-style approach to examining domains rather than syndromes or diseases. On the whole, the intrinsic connectivity findings provide evidence of potential new targets associated with adaptive and maladaptive function, while affirming existing literature, which lends support to the potential neurobiological validity of our data-driven group assignments.

### Limitations

It was not our aim to identify a definitive factor structure from the available behavioral assessments, but rather, to provide a model. Nonetheless, a number of steps were taken to protect against over-or under-fitting the number of factors and provide robust estimates of which subscales loaded on which dimension. Factor number was determined via parallel analysis, factor structure was based on bootstrapping the raw data, and bootstrap-based confidence intervals were computed for estimates of standardized factor loadings. A separate confirmatory sample will be necessary to validate the replicability of the factor structure and the groups identified.

The different response formats of the questionnaires could potentially have lead to clustering of items by comparable response format and hence, recapitulation of the original scale factor structure. There was little evidence of such a result, though. To overcome clustering as a function of similar response formats, all questions would have to be administered with the same response format. However, such an approach would require extremely large samples and known psychometric properties of the scales would be forfeited.

Significant connectome-wide differences decreased dramatically beyond the first level of hierarchical classification. Results at lower levels were only significant with less stringent, RFT multiple comparisons correction (see **Supplementary Information**). There are at least three plausible reasons for this large decrease in significant findings: sample size, group homogeneity, and demographic differences. Average sample size was halved for each incremental level of the hierarchy. Thus, one contributor may merely be decreased power. Another potential factor is increasing subgroup homogeneity: the group factor profiles became progressively more similar to one another at lower levels of the hierarchy. Demographic differences may also contribute to smaller connectome-wide differences at more refined levels of subgroup detection, in that groups may become more demographically similar as they become more behaviorally homogenous. However, cluster analyses were conducted on age and gender regression-residuals, at least mitigating some concerns regarding the influence of these demographic variables.

While biomarker differences may diminish as subgroups become more similar, it is exactly these kinds of comparisons that will ultimately permit differential classification based on differences in neurobiology (57). Larger samples within specific diagnostic categories and/or problem domains will likely provide more power to detect differences among more similar groups (e.g., (18, 19)) and can also remedy some of the problems that arise in exploratory analyses requiring complex multiple comparisons correction. By combining population-based, data-driven categorization and diagnostically focused pattern assessments, we can begin to compare the value of various classification methods. Currently, diagnostic heterogeneity is so marked that only extremely large samples or time-consuming separation into individual diagnostic criteria permit head-to-head comparison of both methods (cf. (58)).

Finally, while the present work focused on phenotypic information alone for group classification/detection, alternative approaches might permit classification based entirely on neurobiology or on a combination of neurobiology and behavioral features. We opted to avoid classification purely on neurobiology as such approaches are complex (59), and interpretation of behavioral characteristics (especially for clinical purposes) can be tricky when groups are clustered entirely on neurobiology. A potentially valuable alternative is analytic approaches that simultaneously consider phenotypic and neurobiological information (60).

### Conclusions

More precise practice in psychiatry is limited by a lack of validated biobehavioral tests and extensive heterogeneity within diagnostic categories. In addition to consensus-based approaches to reframing the current nosology (e.g., RDoC), data-driven approaches to delineating homogenous subgroups, spanning adaptive to maladaptive function can yield clinically meaningful groups with potentially important neurobiological differences. In doing so, it will be important to consider not just symptoms of psychiatric illness (i.e., deficits or problems), but also features of psychiatric health (i.e., strengths or protective factors). Examination of a broad array of phenotypic characteristics, in combination with neurobiological differences may improve our understanding of the pathophysiology of mental illness and provide new preventative and treatment strategies. While these analyses will require large samples and advanced analytic approaches, the present study is evidence that such efforts can yield new hypotheses, as well as support existing theories, helping to focus biological psychiatry on those areas that may yield the highest return on investment.

## Acknowledgments

This work was funded by NIMH R01MH094639, R01MH081218, R01MH083246, R21MH084126, with additional funding for personnel and administrative support provided by grants from the Child Mind Institute, the New York State Office of Mental Health, the Research Foundation for Mental Hygiene, the Brain Research Foundation, and the Stavros Niarchos Foundation. Additional funds were provided by gifts from Phyllis Green, Randolph Cowen, and Joseph P. Healey. All data is publicly available via the International Neuroimaging Data-Sharing Initiative (INDI). Data can be accessed at http://fcon_1000.projects.nitrc.org/ A copyright transfer agreement is required for access to detailed phenotypic information.

### Financial Disclosures

The authors have no biomedical financial interests nor any competing interest to report.

